# An Antibiotic Degrading Engineered Living Material Platform to Combat Environmental Antibiotic Resistance

**DOI:** 10.1101/2023.10.19.563050

**Authors:** Gökçe Özkul, Ebru Şahin Kehribar, Urartu Özgür Şafak Şeker

## Abstract

Presence of antibiotics in natural water bodies is a growing problem regarding the occurrence of antibiotic resistance among various species. This is mainly caused due to excessive use of medical and veterinary antibiotics as well as the lack of effective treatment processes for eliminating residual antibiotic from wastewaters. In this study, we introduce a genetically engineered biomaterial as a solution for effective degradation of one of the dominantly found antibiotics in natural water bodies. Our biomaterial harnesses laccase type enzymes which are known to attack specific type of antibiotics, i.e. fluoroquinolone type synthetic antibiotics, and as a result degradation occurs. The engineered biomaterial is built using *Escherichia coli* biofilm protein CsgA as a scaffold, which is fused separately to two different laccase enzymes with the SpyTag – SpyCatcher peptide-protein duo. The final material is implemented as a part of a living material system. Our proposed system can be integrated as living antibiotic degrading materials to help the combat with antibiotic resistance.

## Introduction

Antibiotics have been the main tool to combatting bacterial infections over the years following the first day of their isolation and production. They have helped, and continue to help the human population survive infections, which otherwise would be lethal, in a comfortable course of disease. In addition to treating infectious disease, discovery of antibiotics made modern, life-saving medical procedures possible, such as organ transplants, and all kinds of surgery [1,2]. Despite their invaluable presence, which have saved millions of lives, nowadays their uncontrolled and irresponsible consumption have trigged a day-by-day growing problem: antibiotic resistance. Antibiotic resistance is an evolutionary process that occurs naturally, however misuse of antibiotics in humans and animals is further accelerating the process. In other words, continuous exposure of the bacteria to antibiotics caused the growth of antibiotic resistance problem. Today, the whole world is facing this inevitable problem and environmental residual antibiotics is an important source of this developing problem [3,4].

Discharges from wastewater treatment plants (WWTP) are one of the two crucial effectors for the release of antibiotics into the environment. The pharmaceutical industry is known to be the examples of critical routes for antibiotics to enter into natural water bodies. These industries produce wastewater containing non-negligible concentrations of antibiotics, which serve as a crucial cause for the occurrence of antibiotics and antibiotic resistance in the nature [5,6]. There are growing number of examples around the world emphasizing the significance of high concentrations of antibiotics found in the pharmaceutical WWTP [7–9]. Another contributor to the occurrence of antibiotics in fresh water bodies is the release of un-processed antibiotics from human and animal sources, which are subjects of wrong and over-consumption of antibiotics. Nearly 50-90 % of the antibiotics used by humans and animals are released from the body as a mixture of original intake antibiotic and its bioactive metabolites in the form of urine and feces [6,10,11].

In the modern wastewater treatment processes, the elimination rates of antibiotics vary significantly on the wastewater characteristics, treatment capacities and systems. It is crucial to point out that, some processes used to treat wastewaters, i.e. biological and disinfection treatments, may even have a deteriorating effect resulting in increasing the variability and concentrations of antibiotics in wastewaters [12,13]. In the current situation, the occurrence and fate of antibiotics during wastewater treatment are not very well understood [14], and the performance of varying treatment processes still need to be documented and analyzed for their success in removing antibiotics from wastewaters [6].

In this current study, we aim to develop an enzymatically active mash structure for treating antibiotics that are found in wastewaters and end up in natural water bodies because they cannot be completely degraded with the existing wastewater treatment processes. Our strategy is to develop a structure based on *Escherichia coli* biofilm protein, CsgA, which is genetically modified to incorporate laccase enzymes. Laccases are robust enzymes, allowing them to function at high temperatures, pH and salt concentrations [15,16]. These features make them strong candidates for using them in wastewaters varying a lot in terms of chemical and physical conditions. We will use two different types of laccase, i.e. CotA [17,18] and YlmD (UniProtKB – O31726). Our strategy shapes around these two enzymes due to their affinity for degrading fluoroquinolone type synthetic antibiotics such as Ciprofloxacin [19]. The mode of action for CotA and YlmD is based on the attacks to the piperazinyl ring of their substrates [20].

Incorporating the aforementioned pieces together, we developed complex biofilm structures composed of CsgA biofilm protein fused with CotA and YlmD, separately (Figure 1). Degradation of Ciprofloxacin with the complex structures and success of this process are analyzed by mass spectroscopy and cell viability assays, respectively (Figure 1).

**Figure 1.**
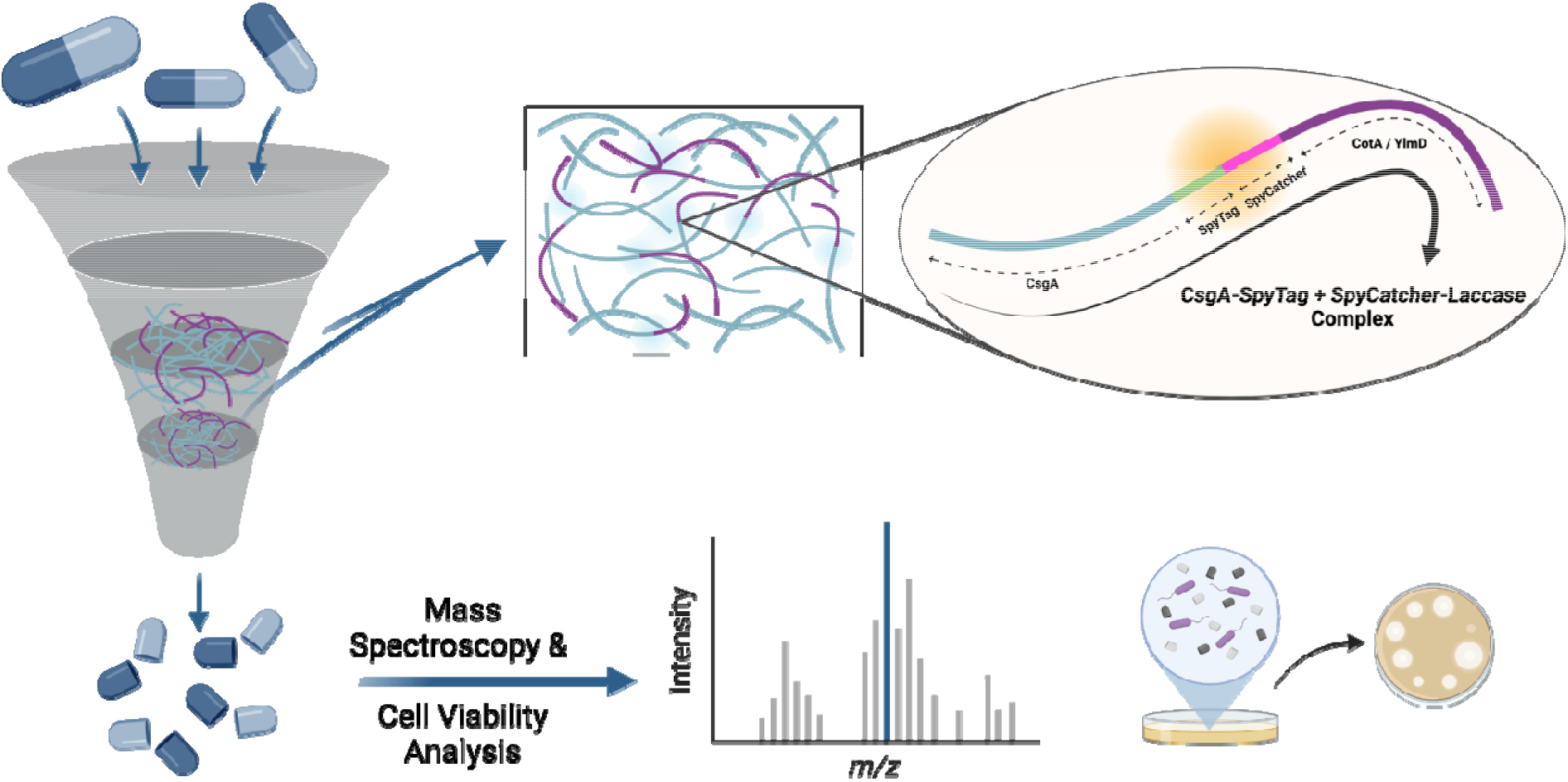
A general interpretation of the functionality of complex structure formed with CsgA-SpyTag + SpyCatcher-Laccases. The formation of CsgA-SpyTag + SpyCatcher-Laccase (i.e. CotA and YlmD) fusion protein. The complex structure is achieved through an irreversible isopeptide bond formation between domains inhabiting SpyTag and SpyCatcher (i.e. CsgA-SpyTag and SpyCatcher-Laccase). Mesh-like structure obtained by the overexpression of CsgA-SpyTag + SpyCatcher-Laccase fusion proteins is used to degrade the residual antibiotic in aquatic environments. This is achieved through the catalytic activity of laccases (CotA and YlmD, separately), which are used to functionalize the mesh structure. End products are analyzed for residual antibiotic activity. Possible degradation products are analyzed using advanced analytical techniques and bacterial cell culture plates.

## Materials and Methods

### Construction of Plasmids for Expressing Complex Subunits

The scaffold of the complex structure, i.e. CsgA-SpyTag, was designed to fuse SpyTag peptide to the C-terminus of CsgA biofilm protein coding gene with the help of a GS linker in between. The SpyCatcher-YlmD and SpyCatcher-CotA constructs were achieved through fusing SpyCatcher protein to the N-terminus of YlmD and CotA proteins, separately along with a GS linker in between the fused protein parts. The gene fragment coding for CsgA-SpyTag fusion protein was cloned into pZa vector under tetO promoter. The fragments coding for SpyCatcher-YlmD and SpyCatcher-CotA fusion proteins were cloned into pET22b (+) vector providing expression under T7 promoter, separately. The plasmid harboring CsgA-SpyT expression cassette was transformed into ΔcsgA strain derived from *E. coli* (MG1655) cells. On the other hand, the plasmids harboring SpyCatcher-YlmD and SpyCatcher-CotA fusion proteins’ expression were transformed into BL21 (DE3) strain.

### Preparation and filtration of CsgA-SpyTag fibers

The preparation of CsgA-SpyTag fibers were conducted as described in our previous paper [21]. Briefly, ΔcsgA cells containing pZa tetO CsgA-SpyTag plasmid were inoculated into lysogeny broth (LB) medium supplied with ampicillin and incubated overnight at 37 °C, 200 rpm. Following, 1:100 dilution was done into fresh LB medium supplemented with ampicillin after the overnight incubation. The cells were grown until OD_600_ reached 0.4. Afterwards, the cells were pelleted and resuspended in 1X M63 minimal medium (pH 7.0, MgSO_4_ (1mM), glycerol (2% v/v), casein hydrolysate (0.1%, v/v), thiamine (1 μg mL^-1^), and ampicillin (30 μg mL^-1^)). The cells were incubated at 30 °C, following stationary conditions. CsgA-SpyTag expression was achieved through aTc (250 μg mL^-1^) addition every two days.

Following induction period, the CsgA-SpyTag fibers were filtered as described earlier [21,34]. Briefly, at the end of incubation period the cell culture was treated with 0.8 M Guanidine-Hydrochloride (Gdn-HCl) for 2 hours at 4 °C. At the end of incubation period, filtration was carried out using a 47 mm polycarbonate filter membrane having 10 μm pores (EMD Millipore). The filter membrane was then incubated with 5 mL of 8M Gdn-HCl for 5 min, vacuumed and washed with 5 mL of dH_2_O, vacuumed. Next, the filter membrane was incubated with 5 mL SDS (5% m/v) for 5 min, vacuumed and washed with 5 mL of dH_2_O, vacuumed. The residual CsgA-SpyTag fibers were then collected via a spatula.

### Preparation of SpyCatcher-CotA and SpyCatcher-YlmD fusion proteins

Cells embracing the plasmids to express SpyCatcher-YlmD and SpyCatcher-CotA fusion proteins were inoculated in fresh LB medium with ampicillin, overnight at 37 °C, 200 rpm. Following incubation, the cells were diluted 1:100 into fresh LB medium supplemented with ampicillin and CuSO_4_ (0.25 mM). The cells containing pET22b SpyCatcher-YlmD plasmid were incubated at 37 °C, 200 rpm and the cells containing pET22b SpyCatcher-CotA plasmid were incubated at 18 °C, 200 rpm until OD_600_ reached 0.4. Afterwards, the cells were induced with IPTG (1 mM) and grown for overnight at their specified temperatures. The cells were pelleted through centrifugation.

Cells were lysed following the resuspension of the pellet in 1X Phosphate Buffer Saline (PBS, pH 7.0). Five rounds of freeze-thaw cycle were applied using liquid nitrogen in the presence of phenylmethyl sulphonyl chloride (PMSF) (1 mM). Sonication took place for 5 min with a 10 second ON and 10 sec OFF cycle with an amplitude of 35%. Afterwards, the sonicated cells were centrifuged at 13000xg for 45 min, and the supernatant was filtered using a 0.20 μm pore size cellulose acetate syringe filter (VWR).

### Complex formation for CsgA-SpyTag + SpyCatcher-YlmD and CsgA-SpyTag + SpyCatcher-CotA

The SpyTag-SpyCatcher complex formation using the CsgA-SpyTag and SpyCatcher-CotA, SpyCatcher-YlmD subunits was conducted as described earlier in our previous work and also by Joshi et al [21,34]. Briefly, CsgA-SpyTag fibers were weighed and aliquoted to have 1 mg/mL final concentration following the induction and filtration processes explained previously. On the other hand, SpyCatcher-YlmD and SpyCacther-CotA fusion proteins were determined to have concentrations of 74.8 μg/mL, 135.7 μg/mL, 220.2 μg/mL (Figure S2, Supporting Information); 26.7 μg/mL, 29.4 μg/mL, and 60.8 μg/mL (Figure S3, Supporting Information), respectively for the complex formation. The complex formation was carried out in Phosphate Buffer (50mM) (1 M KH_2_PO_4_, 1 M K_2_HPO_4_, pH 7.2). The abovementioned subunits were incubated in Phosphate Buffer (50 mM) at 4°C overnight on a shaker. The fiber complexes, i.e. CsgA-SpyTag + SpyCatcher-YlmD and CsgA-SpyTag + SpyCatcher-CotA, were pelleted at 14 000xg for 5 minutes at 10 °C. The supernatant was separated from the fiber complex pellets. Next, the fiber complex pellets were washed 3 times using dH_2_O in order to get rid of the non-specific SpyCatcher-YlmD and SpyCatcher-CotA fusion proteins from their corresponding fiber complex structures.

### Enzymatic Activity Assessment for SpyCatcher-CotA and SpyCatcher-YlmD Fusion Proteins

The enzyme activity of SpyCatcher-YlmD and SpyCatcher-CotA fusion proteins embracing laccase type enzymes, i.e. YlmD and CotA, were assessed through changes in absorbance measurements as a result of 2,2’-azino-bis (3-ethylbenzothiazoline-6-sulfonzuur) (ABTS) substrate degradation [25]. Enzyme activity measurements were conducted after inoculation and cell lysis processes as explained earlier. The enzyme activity for SpyCatcher-YlmD and SpyCatcher-CotA was measured using the whole cell lysate since the yield from Fast Protein Liquid Chromatography (FPLC) was not as high as desired. The enzyme activity assay was conducted as previously described [25]. Briefly, 0.1 M Sodium Acetate Buffer (0.07 M Sodium Acetate, 0.03 M Glacial Acetic Acid, pH 5.0) containing ABTS (5 mM) was mixed with previously defined concentrations of SpyCatcher-YlmD and SpyCatcher-CotA fusion proteins. The absorbance values for indicating ABTS degradation was measured with SpectraMax M5 Microplate Reader (Molecular Devices) spectrophotometer. The absorbance measurements were taken every 20 minutes during 2 hours.

### LCMS-QTOF Analysis

The degradation experiments of Ciprofloxacin were done once the CsgA-SpyTag + SpyCatcher-YlmD and CsgA-SpyTag + SpyCatcher-CotA complexes were formed as described previously. Ciprofloxacin was then added on top the fiber complexes at a concentration of 10 μg/L and incubated at 37°C, for a specified time period. At the end of incubation, the samples were centrifuged at 14000xg for 5 minutes in order to pellet the fiber complex and remove the supernatant for analysis using Liquid Chromatography Mass Spectrometry Quadrupole Time of Flight (LC-MS QTOF). The supernatant was analyzed to check the presence of degradation products which occurred as a result of Ciprofloxacin degradation due to the fiber complexes of CsgA-SpyTag + SpyCatcher-YlmD and CsgA-SpyTag + SpyCatcher-CotA. The samples were transferred into LC-MS QTOF vials and they were analyzed in Agilent LC-MS QTOF 6530. The analysis was conducted without a column and the operating conditions were as following: 300°C for Gas Temperature, 8 L/min for Drying Gas Flow, 35 psi for Nebulizer, 250°C for Sheath Gas Temperature, 6 L/min for Sheath Gas Flow, 3500 V for capillary voltage (Vcap), and 170 V for fragmentor voltage. The analysis was conducted for positive ions.

### Antibiotic Effectivity Analysis for Degradation Products

*E. coli* ΔcsgA cells harboring pZa tetO CsgA-SpyTag plasmid were grown inoculated in fresh LB overnight at 37 ⁰C, 200 rpm. At the end of incubation period, cells were diluted 1:100 into fresh LB medium supplemented with 30 mg/mL Ampicillin and grown until OD_600_ of 0,4 is achieved. Once the desired optical density for cells is achieved, the cells were pelleted at 8000 xg for 5 minutes. Supernatant is removed and the cell pellet is resuspended in fresh 1x M63 minimal media (pH 7.0, MgSO_4_ (1mM), glycerol (2% v/v), casein hydrolysate (0.1%, v/v), thiamine (1 μg/mL^-1^) supplemented with the appropriate antibiotic. The cells were transferred to 24 well plates with a volume of 2 ml per well. The cells are induced to express CsgA-SpyTag with aTc (250 μg/mL) every 2 days and the 1x M63 media is renewed every 3 days without disturbing the cells attached to the bottom of the wells. The total induction period is kept for 8-10 days. Once the induction period is over, M63 media is removed without disturbing the cells at the bottom of wells. Wells were washed with distilled water and the water was removed by quickly inverting the 24 well plate. This washing step was repeated 3 times in order to get rid of the planktonic cells. CsgA-SpyTag +SpyCatcher-CotA and CsgA-SpyTag +SpyCatcher-YlmD complex formation were held within the wells by adding SpyCatcher-CotA and SpyCatcher-YlmD fusion proteins into the desired wells, separately. Following complex formation with using viable cells, antibiotic degradation products were added on top of the complex structures and the cell viability were controlled with optical density measurements.

### Western Blot

The protein samples including SpyCatcher-YlmD and SpyCatcher-CotA were mixed with 1X SDS loading dye (Laemmli Sample Buffer) and incubated at 95 °C for 5 minutes. Complex samples CsgA-SpyTag + SpyCatcher-YlmD and CsgA-SpyTag + SpyCatcher-CotA were incubated with 100% Hexafluoro isopropanol (HFIP) overnight, mixed with 2X SDS loading dye and incubated at 95 °C for 5 minutes.

The protein samples were analyzed with 15% SDS gel. The samples were run through the SDS gel for 2 hours at 120 Volts and then transferred to a polyvinylidene difluoride (PVDF) membrane (Thermo Scientific) via an electroblotting system (BioRad). Following transfer, the membrane was incubated with 5% blocking buffer (non-fat milk powder in Tris Buffered Saline with 0.1% Tween-20, i.e. TBS-T) at 4 °C overnight on a shaker. The PVDF membrane was then incubated with anti-his mouse primary antibody (Thermo Scientific Pierce) in TBS-T (1:5000) for an hour at room temperature on a shaker. The membrane was then washed with TBS-T for 3 times. Following the membrane was incubated with anti-mouse horseradish peroxidase (HRP) conjugated secondary antibody in TBS-T (1:10 000). The membrane was washed again with TBS-T 3 times and the membrane was visualized with Vilbert Lourmat imaging system following an incubation with enhanced chemiluminescence (ECL) substrate (Biorad).

### Statistical Analysis

Data expressed as mean ± SD. For analyzing the significance between data sets, one-way ANOVA testing followed by a Sidak’s multiple comparisons test was carried out across groups. In all cases, significance was defined as *p* ≤ 0.05. Statistical analysis was carried out using GraphPad Prism-6 Software.

## Results and Discussion

### Complex formation with CotA and YlmD Laccases

The formation of CsgA-SpyTag + SpyCatcher-YlmD and CsgA-SpyTag + SpyCatcher-CotA complexes was investigated through analyzing the overexpression of individual subunits, i.e. SpyCatcher-YlmD, SpyCatcher-CotA, and CsgA-SpyT, and the assembled complex structures. CsgA-SpyTag subunit was obtained via expression from aTc inducible plasmid coding for the fusion protein, which is transformed in ΔcsgA cells derived from *E. coli* (MG1655). The CsgA-SpyTag fibers were confirmed using Transmission Electron Microscopy (TEM) (Figure 2). The CsgA-SpyTag biofilm fibers were analyzed using the same methods in our previous work [21]. The fusion fibers were labelled with Ni-NTA conjugated gold nanoparticles, known for their high tendency to interact with Histidine tags [22], which during the design of CsgA-SpyTag fusion protein, we have added to the 3’ end of the fusion protein. In addition, we also verified the bacterial biofilm overexpression using Congo Red (CR) dye, which is known to bind and quantify β-sheet rich amyloid structures [23]. Quantification and formation of CsgA-SpyTag fusion protein was compared with ΔcsgA cells, which essentially lack the gene cassette for CsgA biofilm protein expression (Figure S1, Supporting Information). The SpyCatcher-YlmD and SpyCatcher-CotA subunits were overexpressed using an Isopropyl β-D-1-thiogalactopyranoside (IPTG) inducible pET22b (+) plasmid transformed into *E. coli* BL21 DE3 cells, separately.

**Figure 2.**
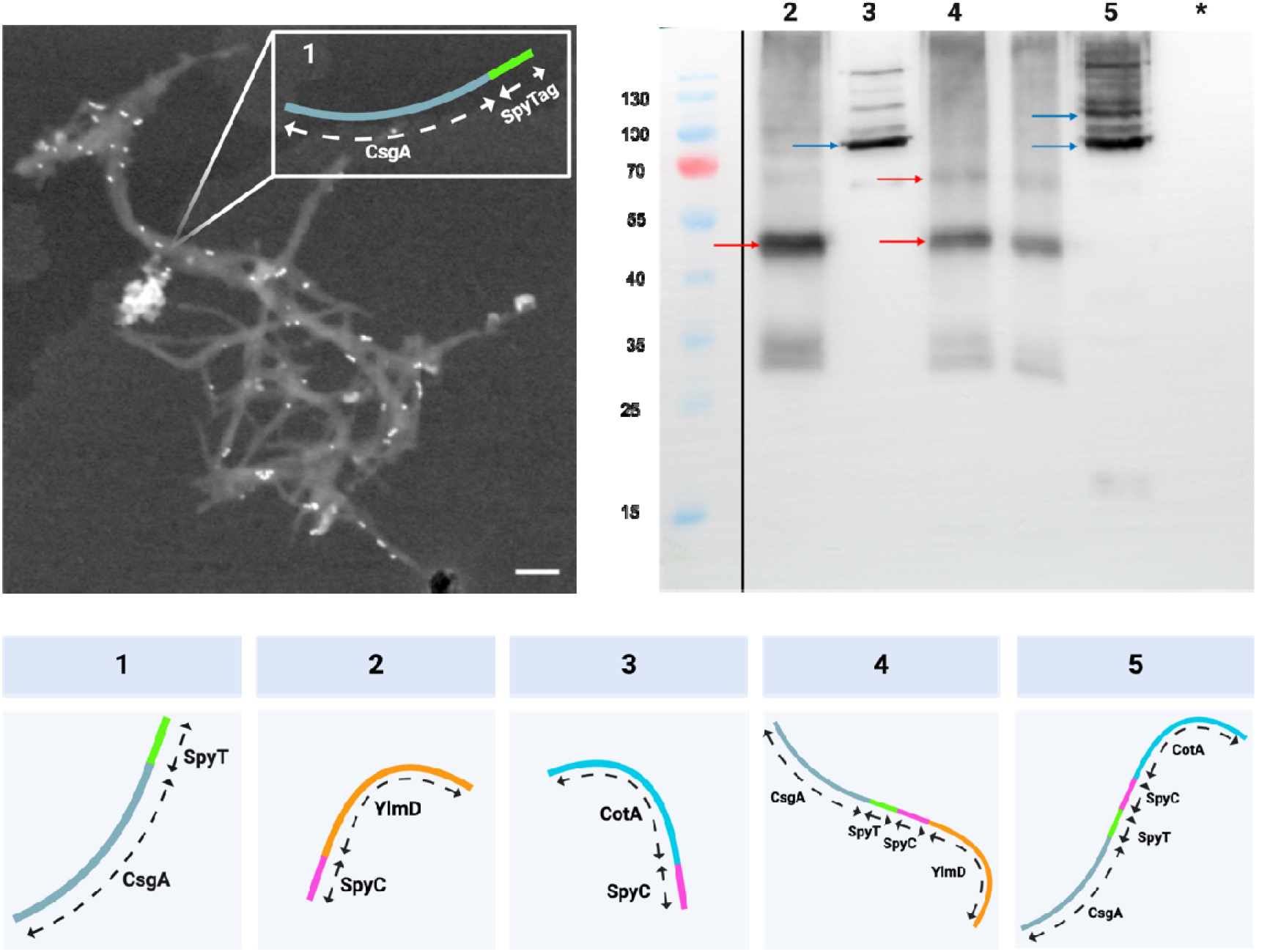
Assessment of individual subunits and complex structures. TEM image of Ni-NTA conjugated Au nanoparticle labelled CsgA-SpyTag fusion protein. The scale bar represents 100 nm. Individual subunits and complex structures imaged via Western Blot. Box 1 indicates CsgA-SpyTag, Box 2 represents SpyCatcher-YlmD, Box 3 represents SpyCatcher-CotA, Box 4 represents CsgA-SpyTag + SpyCatcher-YlmD, Box 5 represents CsgA-SpyTag + SpyCatcher-CotA, the asterisk (*) represents E. coli BL21 DE3. The red arrows indicate SpyC-YlmD and CsgA-SpyTag + SpyCatcher-YlmD complex in Lanes 1 and 3, respectively. The blue arrows indicate SpyC-CotA and CsgA-SpyTag + SpyCatcher-CotA complex in Lanes 2 and 4, respectively. Boxes and the numbers on the TEM and Western Blot images correspond to each other.

The approach for forming complex structures using SpyTag and SpyCatcher subunits is achieved using the same mechanism as explained in our previous work [21]. Briefly, an irreversible isopeptide bond formation is promoted in a favorable reaction medium between the unprotonated amine group of Lys^31^ and carbonyl of Asp^117^ [24]. The verification for the subunits and the complex structures were achieved through Western Blot analysis (Figure 2). SpyCatcher-YlmD, with a 45 kDa molecular weight, and SpyCatcher-CotA, with ∼ 80 kDa, are seen on Lane 2 (red arrow) and 3 (blue arrow), respectively. On Lane 4, The CsgA-SpyTag + SpyCatcher-YlmD complex (60 kDa) is indicated with the upper red arrow, which is formed as a result of the irreversible bond formation between CsgA-SpyTag (15 kDa) and SpyCatcher-YlmD (45 kDa, lower red arrow) subunits [24]. The CsgA-SpyTag + SpyCatcher-CotA (∼ 100 kDa) complex is shown in Lane 5 with the upper blue arrow which is formed as a result of the irreversible interaction between CsgA-SpyTag (15 kDa) and SpyCatcher-CotA (80 kDa, lower red arrow) subunits. The Lanes 4 and 5 accommodate both the complex structures and the subunits forming the complexes, which indicates the complex samples also contain some free SpyCatcher-YlmD and SpyCatcher-CotA subunits.

### Enzymatic activity assessment for SpyCatcher-CotA and SpyCatcher-YlmD

The enzymatic activity of laccase proteins CotA and YlmD can be analyzed through the monitoring absorbance change as 2,2’-azino-bis(3-ethylbenzothiazoline-6-sulfonic acid) (ABTS) is oxidized by the laccase enzymes [25]. The design of SpyCatcher-YlmD and SpyCatcher-CotA fusion proteins was achieved with the aim of not compromising the enzyme activity of the laccase enzymes. Therefore, the enzyme activity assessment for SpyCatcher-YlmD and SpyCatcher-CotA was crucial since they provide the complex structures the capability of laccase activity. The oxidation of phenolic substances, such as ABTS, by laccases is driven by the difference in redox potentials between copper site of the enzyme, specifically T1 copper site, and the corresponding substrate [25].

The catalytic analysis of enzymes revealed that laccase enzymes YlmD and CotA were able to restore their enzymatic activity when they are fused with SpyCatcher domain (Figure 3). However, a significant difference in the oxidizing capacity for SpyCatcher-CotA and SpyCatcher-YlmD was observed when they are compared. For both of the fusion proteins, 3 different concentrations were compared with each other for each fusion protein. For both of SpyCatcher-CotA and SpyCatcher-YlmD, the highest tested concentration showed best enzyme activity. Over a time course of 24 hours, SpyCatcher-CotA was more active enzymatically when compared to SpyCatcher-YlmD and it reached a plateau after 50 minutes for the highest concentration. As a result, we concluded that SpyCatcher-CotA is enzymatically more favorable for use in complex formation. However, we continued to compare both of the laccase fusion proteins for the rest of the experiments.

**Figure 3.**
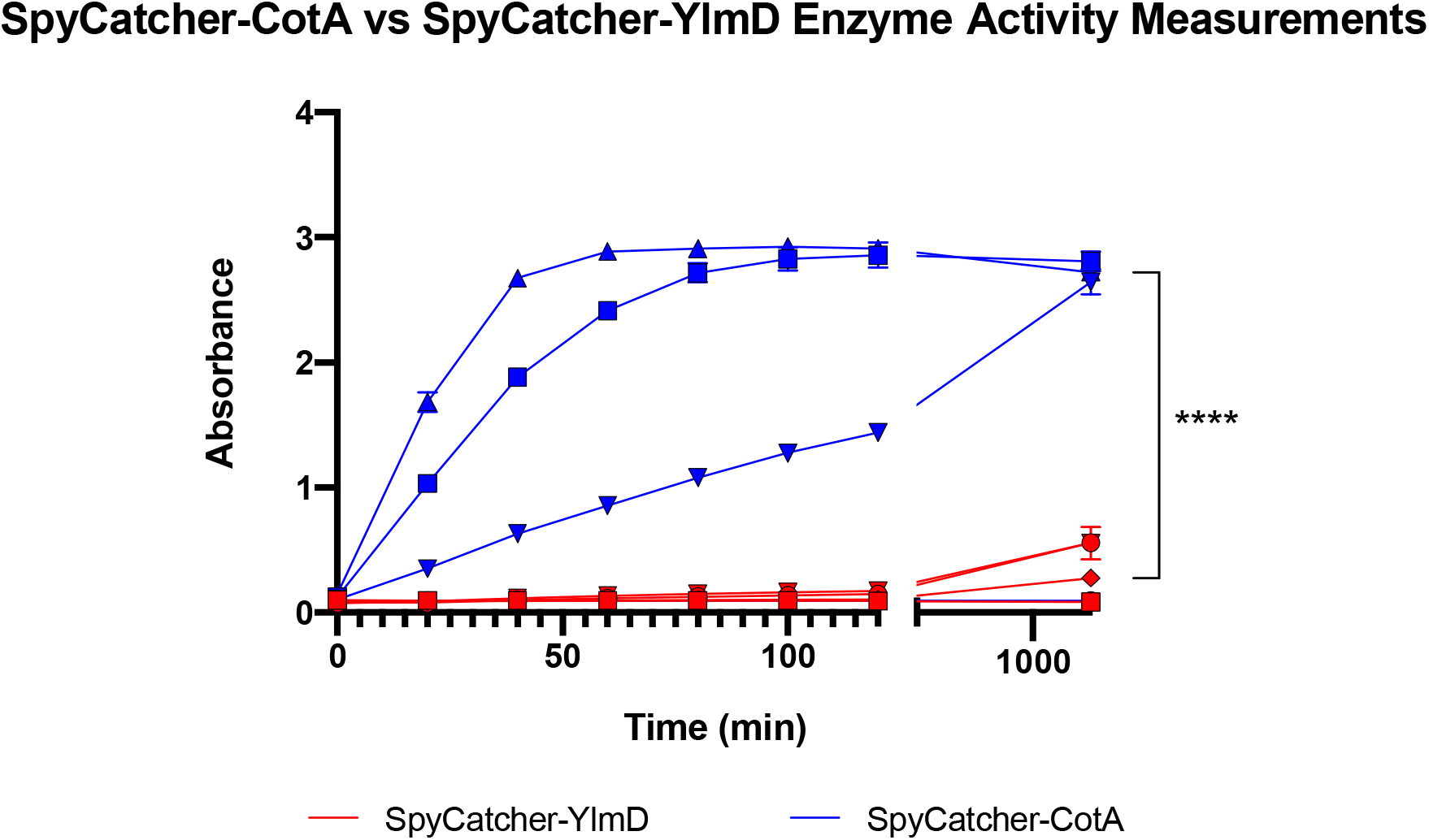
Enzyme activity for catalytic subunits of complex structure (i.e. SpyCatcher-CotA and SpyCatcher-YlmD). Laccase enzyme activity of SpyCatcher-CotA and SpyCatcher-YlmD fusion proteins were analyzed through oxidation of ABTS. As ABTS is oxidized its color changes which is detected by measuring absorbance at 420 nm. The absorbance readings were conducted for 24 hours with measurements at every 20 min for the first 2 hours. For SpyCatcher-YlmD, the red line with circle represents 220,2 μg/mL, red line with tile represents 135,7 μg/mL, and red line with square represents 74,8 μg/mL. For SpyCatcher-CotA, the blue line with triangle represents 60,7 μg/mL, blue line with square represents 45,3 μg/mL, and blue line with inverted triangle represents 26,7 μg/mL.

### Degradation of Ciprofloxacin with CotA and YlmD complexes

Once confirming the enzyme bearing subunit of the complex system is retaining its catalytic activity, we continued with analyzing CsgA-SpyTag + SpyCatcher-YlmD and CsgA-SpyTag + SpyCatcher-CotA complex structures’ ability to degrade the fluoroquinolone type antibiotic, Ciprofloxacin. Laccase type enzymes, i.e. CotA and YlmD, are known to degrade fluoroquinolone type antibiotics through attacking the piperazine ring in the structure of the antibiotic. The degradation products have been previously analyzed with Liquid Chromatography Mass Spectroscopy – Quadrupole Time of Flight (LCMS-QToF) [19,20,26].

Accordingly, CsgA-SpyTag + SpyCatcher-CotA and CsgA-SpyTag + SpyCatcher-YlmD complexes were formed with 4 different concentrations of enzyme subunits each (i.e. 100 μg/mL, 200 μg/mL, 400 μg/mL, 800 μg/mL of SpyCatcher-CotA and SpyCatcher-YlmD). Complexes with varying concentrations of enzyme subunits were tested for their degradation activity against 10 μg/L of Ciprofloxacin. 10 μg/L of Ciprofloxacin is the highest concentration that is found to be present in sewage systems containing discharges from hospitals and urban areas in various cities of Europe and United States [27–30]. Following the incubation of Ciprofloxacin with the formerly mentioned concentrations of CsgA-SpyTag + SpyCatcher-CotA and CsgA-SpyTag + SpyCatcher-YlmD complexes, occurrence of degradation products were analyzed in LCMS-QToF.

The analysis revealed that one specific degradation product (m/z 190) was identified in the samples that were withdrawn from the tubes containing the highest concentration of complex structures of CotA and YlmD laccases, i.e. 800 μg/mL SpyCatcher-CotA and 400 μg/mL, 800 μg/mL SpyCatcher-YlmD, respectively (Figure 4). The degradation product that is encountered in the analysis was initially identified in a previous research focusing on Ciprofloxacin degradation products [26]. This degradation product has a chemical formula of C_10_H_8_NO_3_, which lacks the piperazine ring that is originally present in the Ciprofloxacin [26]. This verifies that the degradation product is obtained with the previously mentioned attack mechanism of laccase type enzymes. When comparing the intensities of the degradation product (m/z 190), CsgA-SpyTag + SpyCatcher-YlmD complex structure is found to degrade Ciprofloxacin when it is formed with 400 μg/mL and 800 μg/mL SpyCatcher-YlmD. However, comparing the intensities of m/z 190 obtained with both of the complex structures, we found the intensities to be almost the same (Figure 4). LCMS-QTOF data can be provided if needed.

**Figure 4.**
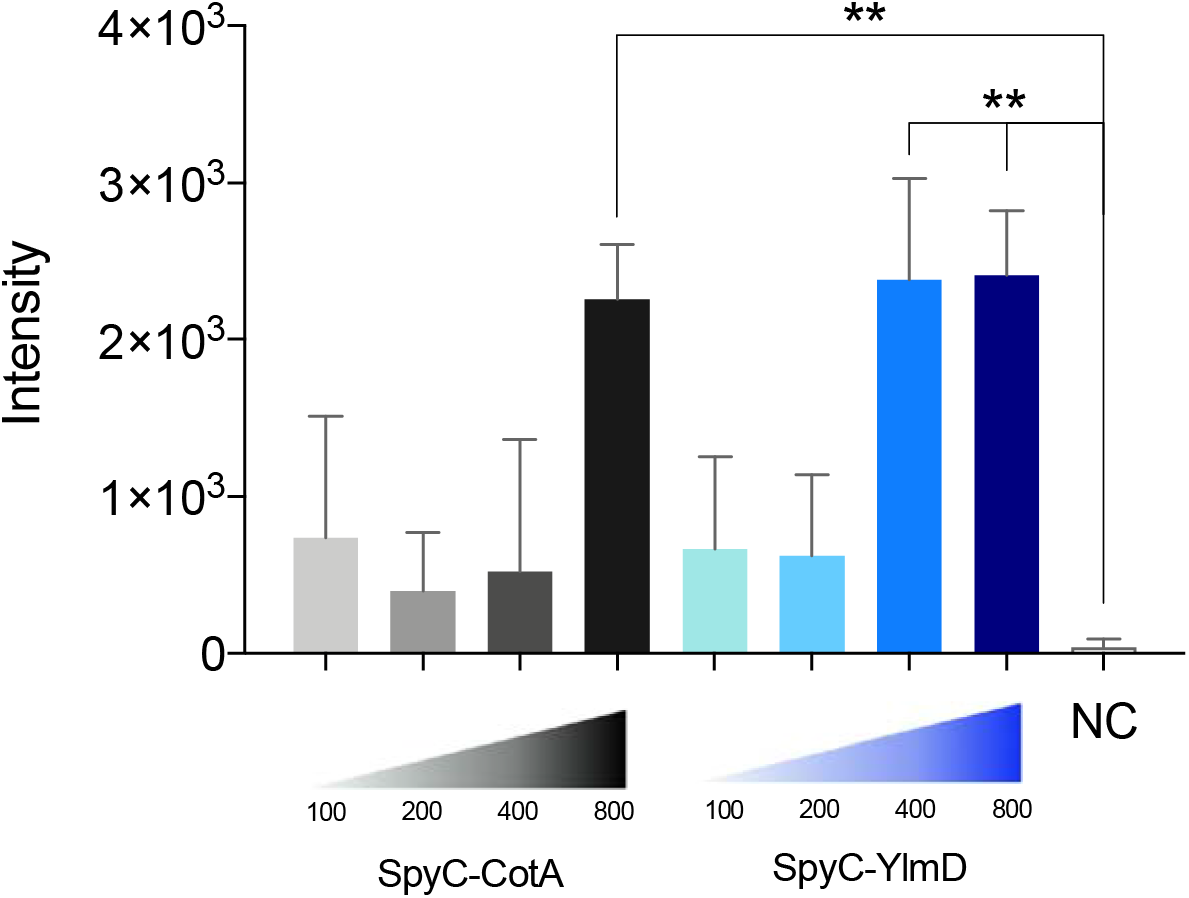
Analysis of Ciprofloxacin degradation products after incubation of Ciprofloxacin with CsgA-SpyTag + SpyCatcher-CotA and CsgA-SpyTag + SpyCatcher-YlmD complexes. The complex structures formed with highest concentrations of SpyCatcher-CotA (black bar) and SpyCatcher-YlmD (blue and dark blue bars) showed significant occurrence of a specific Ciprofloxacin degradation product (m/z 190). The complex structures formed with 800 μg/mL SpyCatcher-CotA, and 400 μg/mL and 800 μg/mL SpyCatcher-YlmD are capable of degrading Ciprofloxacin more than that of the complex structures formed with lower concentrations of SpyCatcher-CotA and SpyCatcher-YlmD. NC represents the negative control, which is CsgA-SpyTag + SpyCatcher complex, which lacks both CotA and YlmD laccases.

### Degradation of ciprofloxacin with cells secreting the complex scaffold

Following the identification of degradation product (m/z 190) as a result of incubating Ciprofloxacin with CsgA-SpyTag + SpyCatcher-CotA complex and CsgA-SpyTag + SpyCatcher-YlmD complex separately, we investigated if *E. coli* MG1655 ΔcsgA cells harboring pZa tetO CsgA-SpyTag plasmid were able to thrive in the presence of Ciprofloxacin that is to be degraded by the complex structures. The cell viability of *E. coli* MG1655 ΔcsgA cells expressing CsgA-SpyTag subunit (with the induction of pZa tetO CsgA-SpyTag plasmid) was crucial in order to make sure the cells forming the base of the complex structure, i.e. CsgA-SpyTag fibers, are able to continuously overexpress CsgA-SpyTag subunit. The CsgA-SpyTag subunit is then used to form the complex structure with the SpyTag – SpyCatcher chemistry.

For this purpose, *E. coli* MG1655 ΔcsgA cells harboring pZa tetO CsgA-SpyTag plasmid were grown and induced with aTc to express CsgA-SpyTag for 8 days in multi-well culture plates. Following the induction period and making sure the cells were successful to express CsgA-SpyTag fibers (Figure S1, Supplementary Information), Ciprofloxacin was introduced to the cell suspension in multi-well plates. A widely used antibiotic, Chloramphenicol, was used to compare the cell viabilities in the presence of antibiotics. Ciprofloxacin was added to the cell suspension in 6 different concentrations (200 mg/mL, 20 mg/mL, 5 mg/mL, 1 mg/mL, 0.1 μg/mL), and Chloramphenicol was added in 3 different concentrations (3400 μg/mL, 340 μg/mL, and 34 μg/mL). Optical density (OD_600_) values of *E. coli* MG1655 ΔcsgA cells expressing CsgA-SpyTag fibers in the presence of Ciprofloxacin showed that CsgA-SpyTag mesh was capable of creating a protected environment for the *E. coli* MG1655 ΔcsgA cells to overexpress CsgA-SpyTag fibers over induction. However, this protected environment can only be maintained if the Ciprofloxacin concentration was kept at 10 mg/mL and lower (Figure 5). Moreover, the tested concentration of Ciprofloxacin for degradation (10 μg /L) is also within the interval where *E. coli* MG1655 ΔcsgA cells can stay viable in the CsgA-SpyTag mesh. As a result, the cells expressing CsgA-SpyTag subunit forming the base of the complex structures can be maintained in the presence of Ciprofloxacin, given a specific concentration range of 0.1 μg /mL and 10 mg/mL is provided.

**Figure 5.**
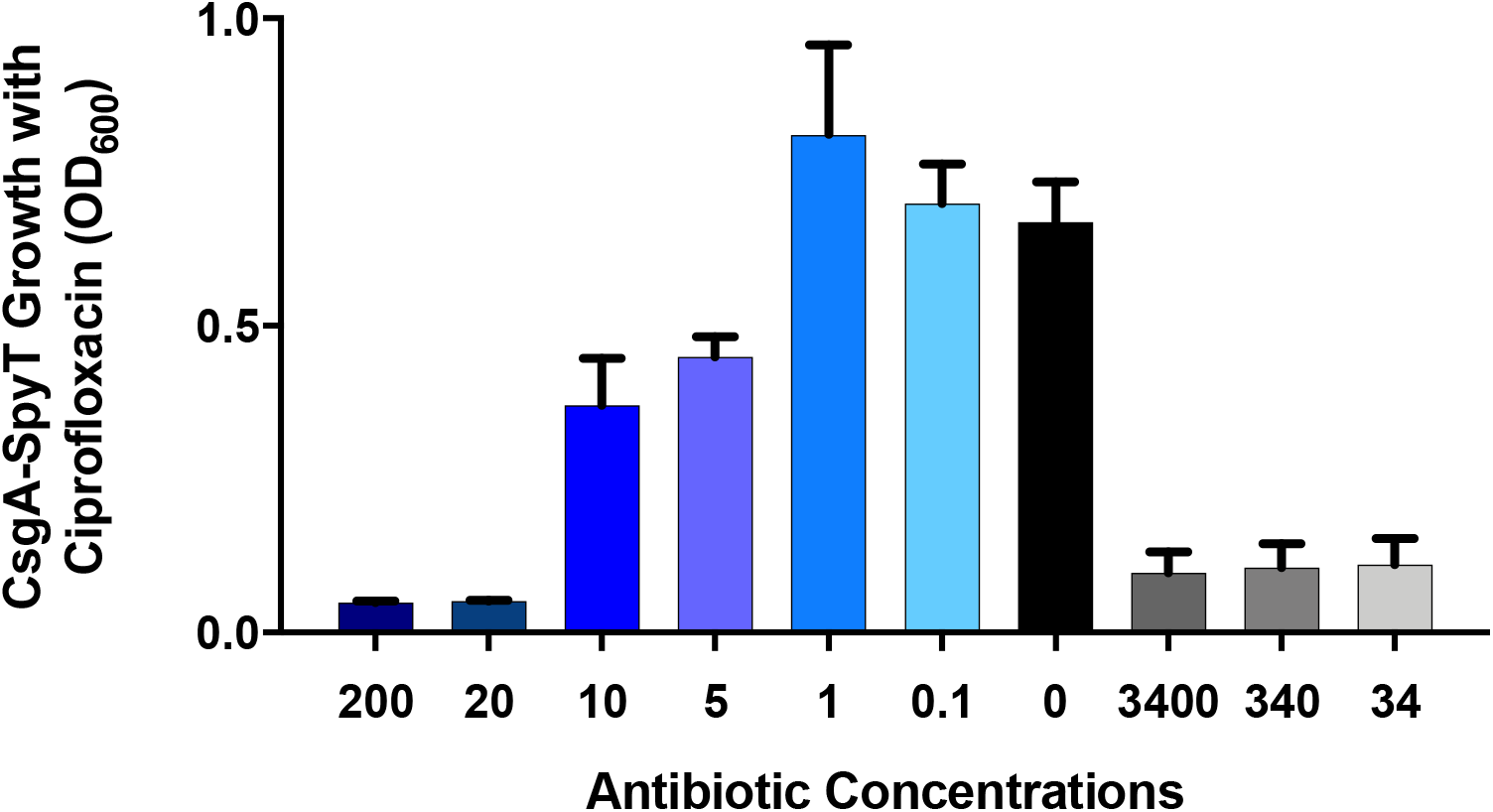
Viability assay for E. coli MG1655 ΔcsgA cells harboring pZa tetO CsgA-SpyTag plasmid in the presence of different concentrations of Ciprofloxacin (200 mg/mL, 20 mg/mL, 5 mg/mL, 1 mg/mL, 0.1 μg/mL, 0 mg/mL, respectively). Chloramphenicol (3400 μg/mL, 340 μg/mL, and 34 μg/mL, respectively) was used as control in order to compare the cell viability in the presence of antibiotics other than Ciprofloxacin. Optical densities of cells in the presence of Ciprofloxacin (between of 0.1 μg /mL and 10 mg/mL) indicated that cells were viable and can continue to express CsgA-SpyTag upon induction with aTc.

### Assessing antibiotic effect of Ciprofloxacin degradation products on E. coli MG1655 cells

Interaction of Ciprofloxacin with the CsgA-SpyTag + SpyCatcher-CotA complex and CsgA-SpyTag + SpyCatcher-YlmD complex, separately resulted in degradation products. It is of crucial importance to identify the antibiotic properties of these products since the ultimate goal is to degrade Ciprofloxacin to a safe-to-discharge version by eliminating its antimicrobial properties. For this, the supernatant obtained as a result of incubating Ciprofloxacin with the complex structures was tested against *E. coli* MG1655 bacteria. The supernatant contains the degradation products of Ciprofloxacin. Therefore, we performed a viability assay in the presence of the supernatant, i.e. Ciprofloxacin degradation products, aiming to verify the reduced antibiotic property of Ciprofloxacin degradation products. We observed that *E. coli* MG1655 bacteria were able to survive in growth media containing Ciprofloxacin degradation products (Figure 6). This indicates the CsgA-SpyTag + SpyCatcher-CotA and CsgA-SpyTag + SpyCatcher-YlmD complexes were successful in degrading Ciprofloxacin.

**Figure 6.**
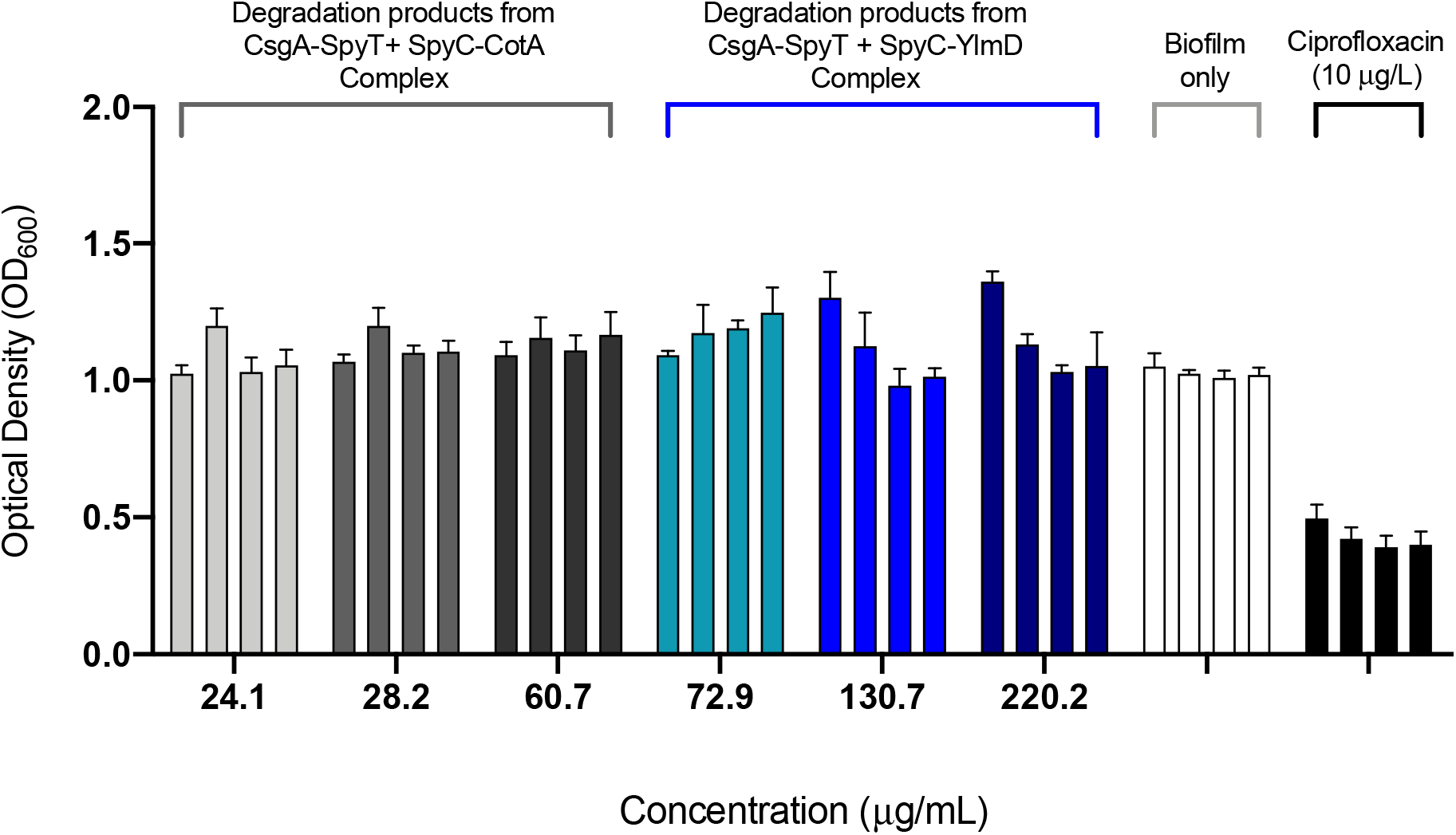
E. coli MG1655 viability assay for Ciprofloxacin degradation products obtained after treatment with CsgA-SpyTag + SpyCatcher-CotA and CsgA-SpyTag + SpyCatcher-YlmD complex structures. The effectivity of antibiotic degradation products is determined according to their antibiotic activity on the cells, hence the optical density analysis. As control group, Chloramphenicol is used and its effect on optical density can be observed.

We have also observed that the viability assay results for cells grown in media containing Ciprofloxacin alone and the degradation products indicate a difference in viability however not significant. We suspect the initial cause can be a mechanism of antimicrobial tolerance in *E. coli* biofilms. The biofilm matrix acts as a physical barrier. This property is addressed to be crucial for limiting and retarding the penetration of antimicrobial agents through the matrix where the cells are embedded [31]. Another property of *E. coli* biofilm structure is the presence of antibiotic degrading enzymes in the matrix. It is found that biofilm matrix includes lyases, transferases, hydrolases and redox enzymes, which can result in antimicrobial tolerance by cleaving or inhibiting the binding of antibiotics [31–33]. The effect of the above-mentioned features of *E. coli* biofilm structures may have a dampening effect on the antimicrobial property of Ciprofloxacin. As a result, we concluded that the *E.* coli MG 1655 cells’ viability assay against Ciprofloxacin degradation products obtained after the incubation period with complex structures harboring the CotA and YlmD laccases, and biofilm matrix without the laccases indicated slight difference (Figure 6; grey, blue and white bars for comparison). The Ciprofloxacin degradation with the presented complex structures, i.e. CsgA-SpyTag + SpyCatcher-CotA and CsgA-SpyTag + SpyCatcher-YlmD, can be enhanced by further modifying the *E. coli* MG1655 cells for reducing its intrinsic property of attacking antimicrobial agents or by further improvement in the activity of biofilm fused CotA and YlmD laccases.

## Conclusion

The presence of antibiotics in natural water bodies is a serious problem for addressing antibiotic resistance. Although the ways antibiotics end up in these systems are known, their removal is still a challenge that needs to be coped with. Therefore, in this study we developed a biofilm based enzymatically active biomaterial that is capable of targeting and destroying a specific type of fluoroquinolone type synthetic antibiotic, Ciprofloxacin. The developed biomaterial is composed of CotA and YlmD fused to *E. coli* biofilm protein, CsgA, separately. The analyzed the complex structure formation for both CsgA-SpyTag + SpyCatcher-YlmD and CsgA-SpyTag + SpyCatcher-CotA, followed by enzyme activity assessment of YlmD and CotA within the complex structure. The complex structures are then tested for Ciprofloxacin degradation. The observed results indicated that the designed complex structures for CsgA-SpyTag + SpyCatcher-YlmD and CsgA-SpyTag + SpyCatcher-CotA, were capable of degrading Ciprofloxacin. In the future, complex systems can be further engineered for enhanced enzyme activity and therefore improved antibiotic degradation. Moreover, the complex structures can be manipulated for serving other purposes such as degradation of hard to degrade chemicals found in natural systems and also wastewaters.

## ACKNOWLEDGEMENTS

We thank to TUBITAK for partial support from the Grant number 119M037. We thank Recep Erdem Ahana (MSc) for fruitful discussion on some of the experimental parts of the manuscript.

## AUTHOR INFORMATION

## SUPPORTING INFORMATION

**Figure S1.**
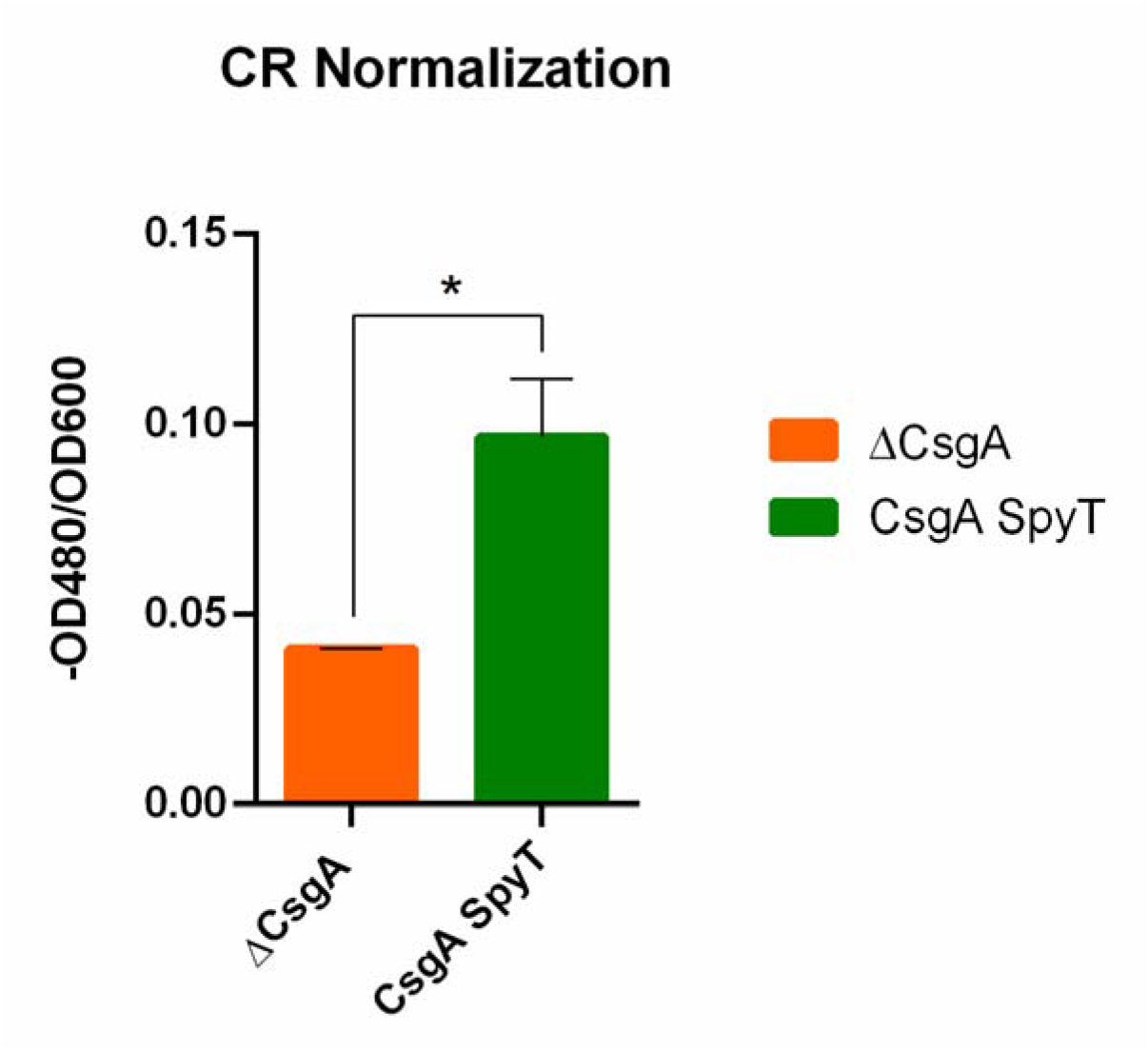
Congo Red Staining for CsgA-SpyTag fibers grown for 3 days. Data is presented as mean ± SD, n=3, *P*-values are calculated using one-way ANOVA with Sidak‘s correction, \**P*<0.05, \*\**P*<0.01.

**Figure S2.**
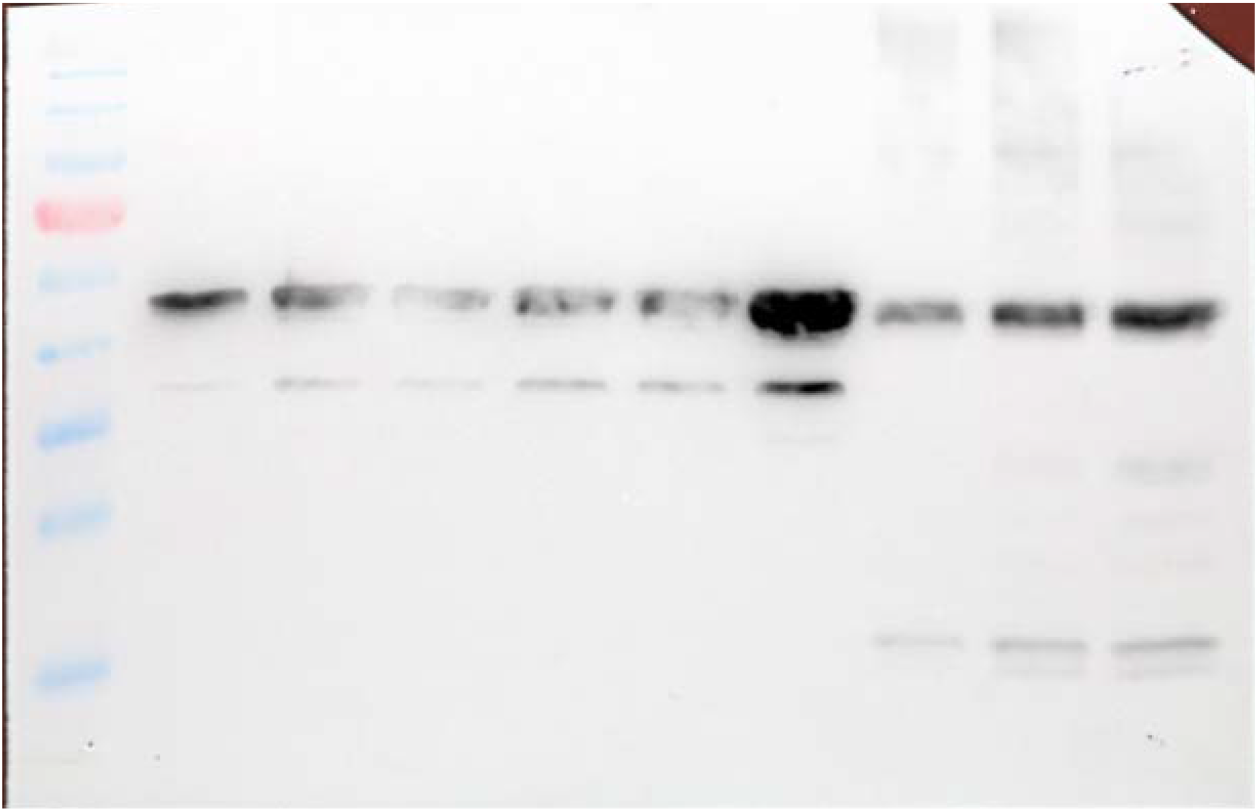
Concentration determination for SpyCatcher-YlmD via Western Blot. Lane 1 represents Page Ruler ladder. Lanes 2, 3, 4, 5, 6, and 7 are 10 μg/mL, 20 μg/mL, 50 μg/mL, 100 μg/mL, 250 μg/mL, and 500 μg/mL of reference protein, respectively. These lanes are used to create a standard curve. Lanes 8, 9, 10 are SpyCatcher-YlmD fusion protein that was used in the experiments. Comparing the intensities of SpyCatcher-YlmD in lanes 8, 9, and 10 with the standard curve created with the reference protein of known concentration in Fusion quantification analysis software of Vilber Lourmat Fusion Solo S imaging system, the concentrations of SpyCatcher-YlmD fusion protein were determined. The calculated concentrations for SpyCatcher-YlmD on lanes 8, 9, and 10 are 74.8 μg/mL, 135.7 μg/mL, and 220.2 μg/mL, respectively.

**Figure S3.**
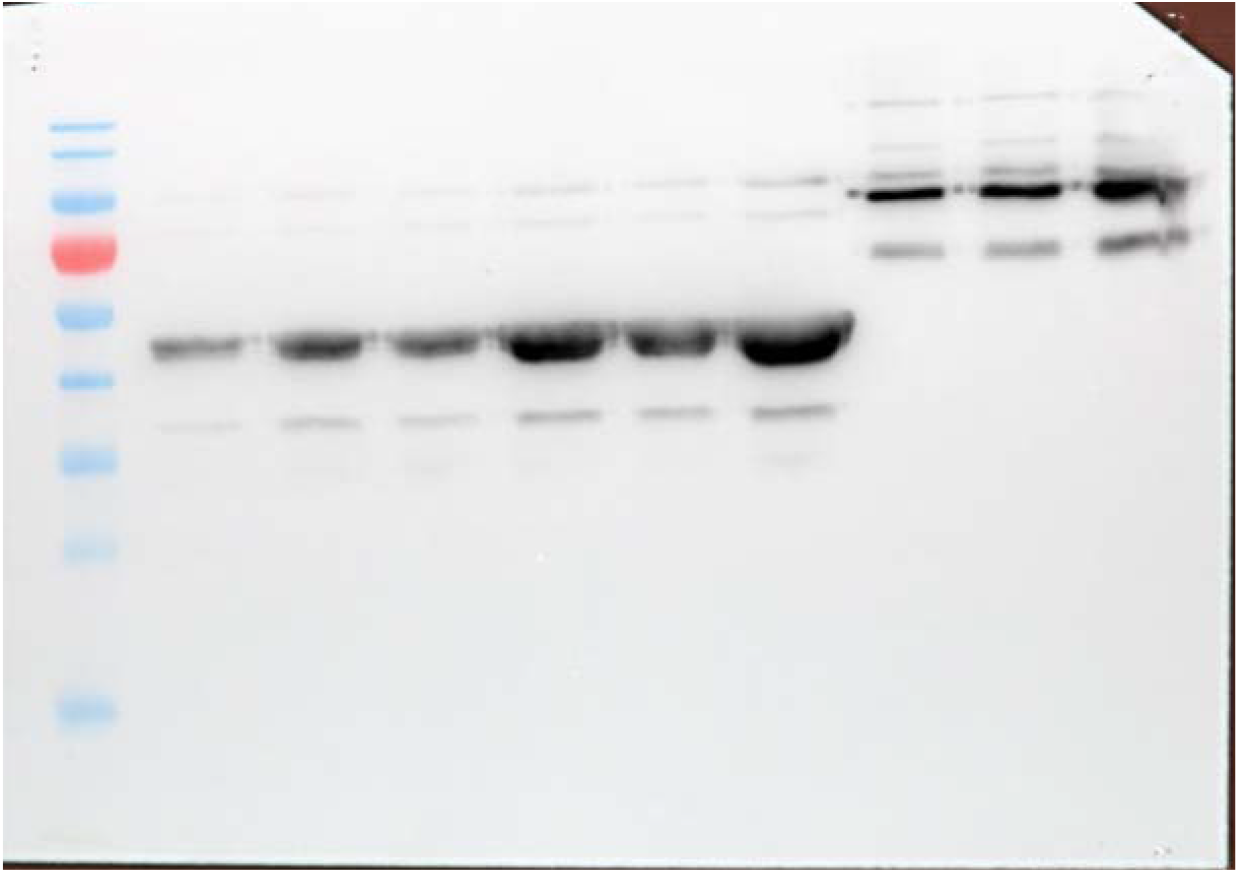
Concentration determination for SpyCatcher-CotA via Western Blot. Lane 1 represents Page Ruler ladder. Lanes 2, 3, 4, 5, 6 and 7 are 10 μg/mL, 20 μg/mL, 50 μg/mL, 100 μg/mL, 250 μg/mL, and 500 μg/mL of reference protein, respectively. These lanes are used to create a standard curve. Lanes 8, 9, 10 are SpyCatcher-CotA fusion protein that was used in the experiments. Comparing the intensities of SpyCatcher-CotA in lanes 8, 9, and 10 with the standard curve created with the reference protein of known concentration in Fusion quantification analysis software of Vilber Lourmat Fusion Solo S imaging system, the concentrations of SpyCatcher-CotA fusion protein were determined. The calculated concentrations for SpyCatcher-CotA on lanes 8, 9, and 10 are 26.7 μg/mL, 29.4 μg/mL, and 60.8 μg/mL, respectively.

## REFERENCES

[1] M.I. Hutchings, A.W. Truman, B. Wilkinson, Current Opinion in Microbiology 51 (2019) 72–80.

[2] World Health Organization, (2020).

[3] RIKILT - BU Toxicology Bioassays & Novel Foods, VLAG, RIKILT - Business unit Dierbehandelingsmiddelen, M.G. Pikkemaat, H. Yassin, H.J. Fels-Klerx, B.J.A. Berendsen, Antibiotic Residues and Resistance in the Environment, RIKILT Wageningen UR, Wageningen, 2016.

[4] S.I. Polianciuc, A.E. Gurzău, B. Kiss, M.G. Ltefan, F. Loghin, Medicine and Pharmacy Reports (2020).

[5] O. Cardoso, J.-M. Porcher, W. Sanchez, Chemosphere 115 (2014) 20–30.

[6] K. Wang, T. Zhuang, Z. Su, M. Chi, H. Wang, Science of The Total Environment 788 (2021) 147811.

[7] H.Q. Anh, T.P.Q. Le, N. Da Le, X.X. Lu, T.T. Duong, J. Garnier, E. Rochelle-Newall, S. Zhang, N.-H. Oh, C. Oeurng, C. Ekkawatpanit, T.D. Nguyen, Q.T. Nguyen, T.D. Nguyen, T.N. Nguyen, T.L. Tran, T. Kunisue, R. Tanoue, S. Takahashi, T.B. Minh, H.T. Le, T.N.M. Pham, T.A.H. Nguyen, Science of The Total Environment 764 (2021) 142865.

[8] P. Kairigo, E. Ngumba, L.-R. Sundberg, A. Gachanja, T. Tuhkanen, Science of The Total Environment 720 (2020) 137580.

[9] X. Liu, S. Lu, W. Guo, B. Xi, W. Wang, Science of The Total Environment 627 (2018) 1195–1208.

[10] J. Menz, O. Olsson, K. Kümmerer, Journal of Hazardous Materials 379 (2019) 120807.

[11] D. Cheng, H.H. Ngo, W. Guo, S.W. Chang, D.D. Nguyen, Y. Liu, Q. Wei, D. Wei, Journal of Hazardous Materials 387 (2020) 121682.

[12] J. Bengtsson-Palme, M. Milakovic, H. Švecová, M. Ganjto, V. Jonsson, R. Grabic, N. Udikovic-Kolic, Water Research 162 (2019) 437–445.

[13] D. Cheng, H. Hao Ngo, W. Guo, S. Wang Chang, D. Duc Nguyen, Y. Liu, X. Zhang, X. Shan, Y. Liu, Bioresource Technology 299 (2020) 122654.

[14] A. Fiorentino, A. Di Cesare, E.M. Eckert, L. Rizzo, D. Fontaneto, Y. Yang, G. Corno, Science of The Total Environment 646 (2019) 1204–1210.

[15] W. Du, C. Sun, J. Liang, Y. Han, J. Yu, Z. Liang, Journal of Food Biochemistry 39 (2015) 101–108.

[16] T.-N. Wang, M. Zhao, Appl Microbiol Biotechnol 101 (2017) 685–696.

[17] P.S. Chauhan, B. Goradia, A. Saxena, 3 Biotech 7 (2017) 323.

[18] F.J. Enguita, L.O. Martins, A.O. Henriques, M.A. Carrondo, Journal of Biological Chemistry 278 (2003) 19416–19425.

[19] A. Prieto, M. Möder, R. Rodil, L. Adrian, E. Marco-Urrea, Bioresource Technology 102 (2011) 10987–10995.

[20] M. Čvančarová, M. Moeder, A. Filipová, T. Cajthaml, Chemosphere 136 (2015) 311–320.

[21] G. Özkul, E.Ş. Kehribar, R.E. Ahan, İ.Ç. Köksaldı, A. Özkul, B. Dinç, S. Aydoğan, U.Ö.Ş. Şeker, Adv Materials Inter 9 (2022) 2201126.

[22] J.A. Bornhorst, J.J. Falke, in: Methods in Enzymology, Elsevier, 2000, pp. 245–254.

[23] C. Reichhardt, A.N. Jacobson, M.C. Maher, J. Uang, O.A. McCrate, M. Eckart, L. Cegelski, PLoS ONE 10 (2015) e0140388.

[24] B. Zakeri, J.O. Fierer, E. Celik, E.C. Chittock, U. Schwarz-Linek, V.T. Moy, M. Howarth, Proceedings of the National Academy of Sciences 109 (2012) E690–E697.

[25] R. Chandra, P. Chowdhary, Environ. Sci.: Processes Impacts 17 (2015) 326–342.

[26] L. Pan, J. Li, C. Li, X. Tang, G. Yu, Y. Wang, Journal of Hazardous Materials 343 (2018) 59–67.

[27] S. Rodriguez-Mozaz, I. Vaz-Moreira, S. Varela Della Giustina, M. Llorca, D. Barceló, S. Schubert, T.U. Berendonk, I. Michael-Kordatou, D. Fatta-Kassinos, J.L. Martinez, C. Elpers, I. Henriques, T. Jaeger, T. Schwartz, E. Paulshus, K. O’Sullivan, K.M.M. Pärnänen, M. Virta, T.T. Do, F. Walsh, C.M. Manaia, Environment International 140 (2020) 105733.

[28] L. Lien, N. Hoa, N. Chuc, N. Thoa, H. Phuc, V. Diwan, N. Dat, A. Tamhankar, C. Lundborg, IJERPH 13 (2016) 588.

[29] M. Gros, S. Rodríguez-Mozaz, D. Barceló, Journal of Chromatography A 1292 (2013) 173–188.

[30] S. Aydin, M.E. Aydin, A. Ulvi, H. Kilic, Environ Sci Pollut Res 26 (2019) 544–558.

[31] V. Ballén, V. Cepas, C. Ratia, Y. Gabasa, S.M. Soto, Microorganisms 10 (2022) 1103.

[32] M.A. Rather, K. Gupta, M. Mandal, Braz J Microbiol 52 (2021) 1701–1718.

[33] C.W. Hall, T.-F. Mah, FEMS Microbiology Reviews 41 (2017) 276–301.

[34] N.-M. Dorval Courchesne, A. Duraj-Thatte, P.K.R. Tay, P.Q. Nguyen, N.S. Joshi, ACS Biomater. Sci. Eng. 3 (2017) 733–741.

[35] L. Pan, J. Li, C. Li, X. Tang, G. Yu, Y. Wang, Journal of Hazardous Materials 343 (2018) 59–67.

